# Parallel Belief States account for Learning and Updating of Attentional Priorities in Multidimensional Environments

**DOI:** 10.1101/2025.09.03.674042

**Authors:** Paul Tiesinga, Thilo Womelsdorf

## Abstract

Inferring the behavioral relevance of visual features is difficult in multidimensional environments as many features could be important. One solution could involve tracking the experience with multiple features and using attentional control to decide which subset of features to explore and chose. Here, we characterize this attentional control process with a model of parallel belief states and test it with a task requiring the learning and updating of attention to features with varying selection histories and motivational costs. We found that exploring and exploiting features was accounted for by a model that tracks the latent beliefs about the relevance of multiple features in parallel. These parallel belief states accounted for the fast learning of feature-based attention, for perseverative selection history effects for features that were previously relevant, and for enhanced learning performance when the motivational costs of making errors increased. Taken together, these results quantify how multiple parallel belief states guide exploration and exploitation of feature-based attention during learning. We suggest parallel belief states represent attentional priorities that are read out by a competitive attentional control process to explore and exploit those visual objects in multidimensional environments that are believed to be relevant.

**Significance statement:** During goal-directed behavior attention is allocated to features relevant for the behavioral goal. But in real-world settings with multidimensional objects, it is often unknown which features are maximally relevant and should be attentionally prioritized. We found a solution to this problem by quantifying the hidden beliefs about the relevance of multiple features in parallel. By tracking belief states about feature relevance we found that subjects consider multiple features in parallel during the learning of feature-based attention. These belief states correspond to attentional priorities and explained when attention is biased towards previously relevant but now irrelevant features, and when learning about relevant features is enhanced by motivational incentives. These findings quantify the parallel hidden beliefs that guide attention in complex environments.

## Introduction

Goal-directed behavior depends on reliable estimates about the relevance of available objects for reaching the behavioral goal. Knowledge of the relevance of objects is, however, challenging when objects are new, vary their relevance over time, or are composed of multiple feature dimensions with possibly conflicting relevance. To make adaptive choices in such multidimensional and dynamically varying environments, subjects benefit from inferring the relevance of multiple features in parallel and weighting their relevance to determine which object to choose. Inferring and representing the relevance of multiple objects differs fundamentally from representing only a few features and calls upon dedicated strategies at the level of neuronal networks, cognitive representations, and with regard to learning mechanisms (Womelsdorf *et al*., 2022; Goudar *et al*., 2023; Johnston & Fusi, 2023).

In neuronal networks, distinguishing two simple stimuli is achievable by representing each in a low-dimensional space and quickly (in a few trials) aligning the decision criterion to this space (Goudar *et al*., 2023) (**Fig. 1A**), while learning to distinguish multiple features requires separate dimensions for each feature and disentangling them in abstract space (Johnston & Fusi, 2023) (**Fig. 1B,C**). In cognitive behavioral studies learning about multiple objects involves representing many features of these objects in parallel (Kriegeskorte & Kievit, 2013; Souza & Oberauer, 2016; Allen & Ueno, 2018; Hitch *et al*., 2018; Cowan *et al*., 2024). On one hand, subjects explore objects whose relevance is uncertain and not yet known, while simultaneously exploit objects whose values have become already predictable (Pearson & Le Pelley, 2020; Rusz *et al*., 2020; Pearson *et al*., 2024). Exploring uncertain options and exploiting predictive features suggest that adaptive choices in multidimensional envrinoments originate from different types of representations that consider multiple objects and their features in parallel. How these parallel representations of the relevance of multiple features are learned with experience has been studied using diverse modeling architectures. Bayesian approaches track the reward value of each object constellation and calculate the optimal weights for attending those objects based on an exhaustive memory of previous rewards and punishments (Leong *et al*., 2017; Oemisch *et al*., 2019). While this approach successfully models human and nonhuman primate learning in some value learning tasks (Leong *et al*., 2017; Oemisch *et al*., 2019), it is computationally demanding and does not generalize to other multitdimensional learning tasks (Womelsdorf *et al*., 2022; Treuting *et al*., 2025). RL models have resolved learning values in complex feature spaces with different strategies. In one account learning in multidimensional environments simplifies the problem space using hypothesis testing instead of inference by testing with each choice the relevance of individual features and switches the hypothesis when choosing the inferred feature results in a negative outcomes (Farashahi *et al*., 2017). Recent RL agents were shown to cope with a large feature space by augmenting a slow RL learner with other cognitive strategies such as working memory (Collins & Frank, 2012), adaptive exploration (Khamassi *et al*., 2011; Treuting *et al*., 2025) and structure learning (Wang *et al*., 2018; Whittington *et al*., 2022). These model frameworks implement versatile mechanisms of learning, but it has remained open which set of mechanisms will generalize across multidimensional tasks and show learning dynamics that match the behavioral time timescales shown by subjects.

**Fig. 1.**
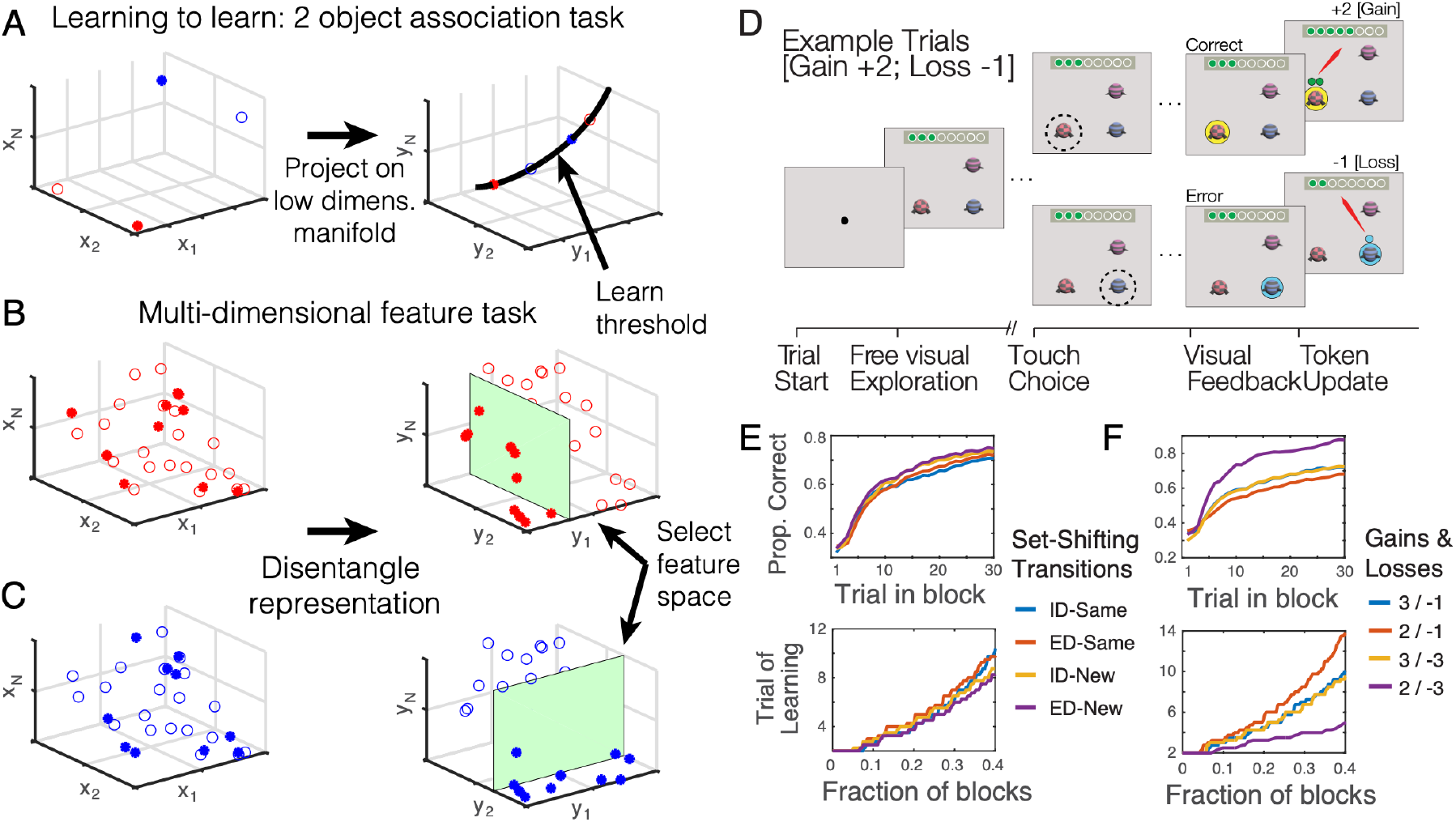
Multi-dimensional learning task requires disentangled representations for attentional control to act on. (**A**) Learning to map 1 of 2 features (e.g. red vs blue) with an action (action 1: open; 2: solid) can be solved by projecting the object space onto a low-dimensional manifold and setting the right threshold. (**B, C**) In a multi-dimensional world objects have many possible features (symbols) of which a subset (e.g. 10) are visible (solid). Learning to map features of the red and blue objects with an action corresponds to dividing objects in two groups, those that contain the target (solid), and those that do not (open). There is unique geometry of the target group (*left*) that can be disentangled (*right*) by mapping each feature to a separate subspace. Learning the green classification plane benefits from attentional control. (**D**) Progression of events in an example trial of the multidimensional learning task with a correct (*upper*) and incorrect (*lower*) outcome. (**E**) Average accuracy (upper) and trial-to-reach learning criterion (bottom) across 4 monkeys. Learning is slightly faster when a block used features that were new (yellow and purple) rather than the same as in the preceding block of trials. (**F**) Performance is better (*upper*) and learning faster (*bottom*) when subjects anticipated loosing 3 tokens for erroneous and gaining 2 tokens for correct choices, which is least motivational challenging condition.

In the absence of a generalizable framework for adaptive learning and decision making in multidimensional environments, we propose an observational model approach that tracks the explicit *belief states* about the relevance of all features subjects encountered during the learning process. The core assumption of such a *Parallel Belief States* (PBS) model is that exploration and exploitation of objects originate from latent states that integrate previous experiences to determine whether objects should be explored, exploited, or avoided (Goudar *et al*., 2024). We show that the transitioning of these latent belief states quantifies which of multiple features is explored because of ongoing uncertainty about their probability to influence object choice, and which features are exploited because of high reward predictability. We apply the PBS model to account for the choices of four nonhuman primates performing a learning and updating task, that requires trial-and-error learning of rewarded target features in the presence of multiple distracting features. The task varies in whether current targets and distractors are novel or used previously, and whether choices have high or low motivational costs. We found that modeling parallel belief states accounted for subjects choices of this multidimensional attentional learning task and accounts for the biases of subjects to explore object features longer when they have a history of being target features and to exploit objects sooner when motivational incentives are higher.

## Results

Four subjects (M1 to M4) performed a feature-based attentional learning task in 44-46 experimental sessions. In each session 36 different target features were learned in blocks of 32-65 trials. Each trial presented three objects, of which the subject chose one (**Fig. 1D**). When the subject chose the one containing the target feature it was rewarded with a gain in number of tokens, whereas when it chose another object it lost a specific number of tokens. The target feature remained the same within a block and changed when a new block started. Blocks differed in three ways. First, the number of tokens gained with a correct choice (+2 or +3) and the number lost after an incorrect choice (-1 or -3) defined the motivational context for each block with four gain/loss combinations: 2/-1, 3/-1, 3/-3,2/-3. Second, blocks differed whether the feature dimension of the target feature remained the same as in the previous blocks (intradimensional block transition, ID), or if the dimension was different (extradimensional transition, ED). Thirdly, in each block the set of feature dimensions present in the objects either stayed the same as in the previous block (Same), or were novel (New). We refer to the feature-based block conditions as New-ID, New-ED, Same-ID, and Same-ED.

The feature-based conditions and the motivational contexts affected how fast subjects learned target features within blocks. Learning occured earlier and performance reached a higher plateau accuracy when blocks transitioned to novel objects rather than reassigned values of previously used objects (New-ID and New-ED versus Same-ID and Same-ED (**Fig. 1E**). In addition, learning was faster when blocks entailed high symbolic punishment (high loss) for erroneous choices (**Fig. 1F**).

### Learning dynamics is reflected in state changes of feature-channels

Learning which feature is a rewarded target requires considering multiple possible features. To formally evaluate how many features subjects consider in parallel during learning we considered individual features as independent channels and quantified the probability that a feature channel represents relevant target or non-target using a Hidden Markov Model (HMM). The HMM used the previous trial outcome and the previously chosen features to update the hidden state of a feature in the current trial (Bengio & Frasconi, 1994) (**Fig. 2A**). The state of a feature channel corresponds to the belief that the feature is relevant. Updating this belief takes into account whether the feature was chosen or not (C+, C-) and whether it was rewarded or not (R+,R-). We found that the choices of subjects were best modelled with four distinct hidden belief states that gradually changed during learning (**Suppl. Fig. 1A,B**). The hidden states correspond to the uncertainty of believing that a feature is relevant, which ranges from being certain that it is relevant (p>0.9, Persist state) or that it is not relevant (p<0.3, Avoid state), to being uncertain about it relevance (p=∼0.3, Explore state), to a somewhat certain belief that a feature is relevant (p>∼0.4-0.8, Preferred state) (*see* Methods). Similar state definitions have been used to account for rule-based switching behavior (Goudar *et al*., 2024). The four states systematically vary over trials and closely follow the learning progress. Example traces of the feature specific belief states showed that when a feature is not the target feature its feature channel transitioned within a few trials to the Avoid state, while when it becomes the target feature it transitions through the Explore- and Preferred-state, reaching the Persist state when learning completed (**Fig. 2B**).

**Fig. 2.**
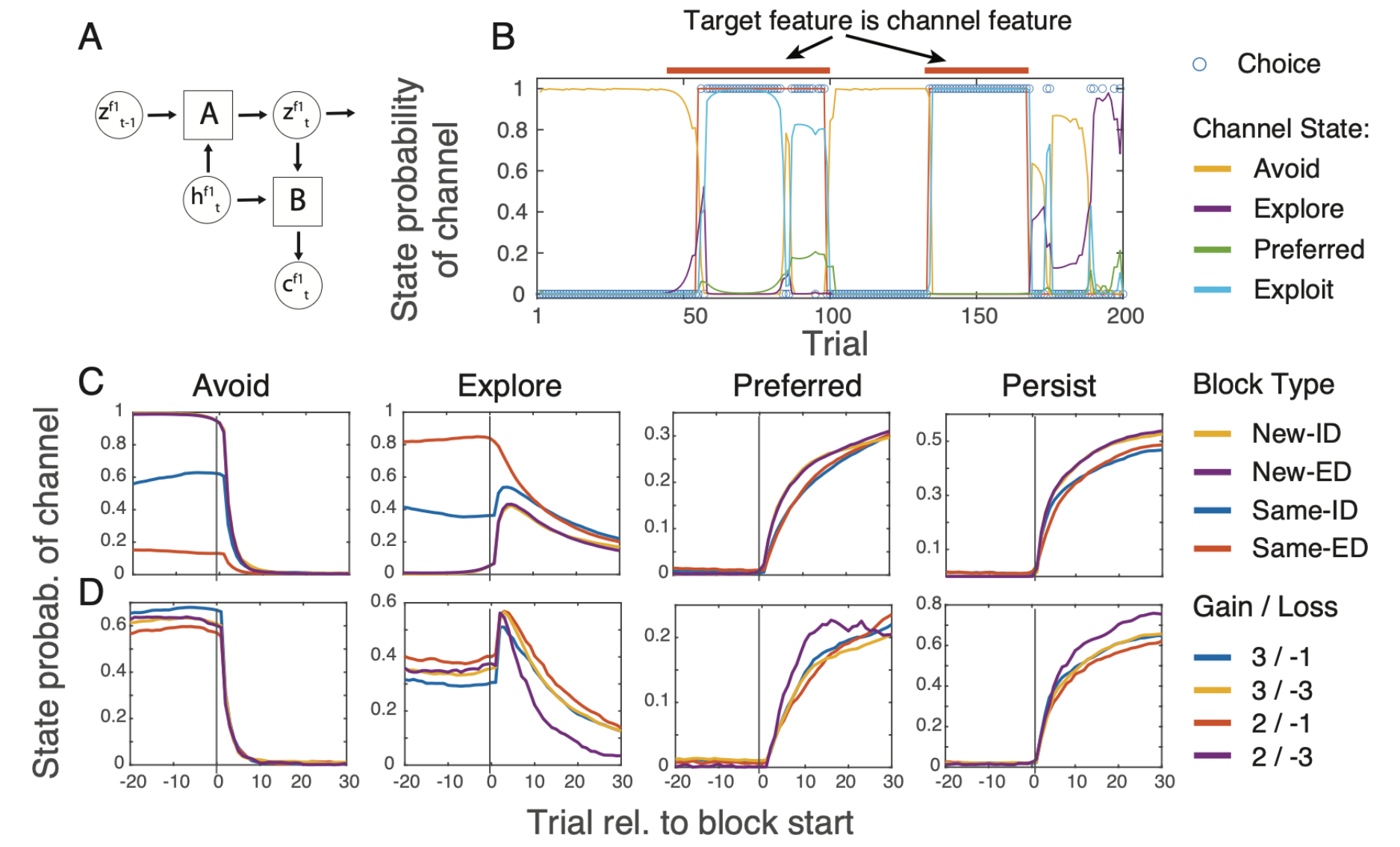
Model overview and behavioral fits showing faster transitions of the target feature state to the Preferred and Persist states when target features are novel and anticipated losses are large. (**A**) Overview of model structure, indicating the predicted choice *c*_*t*_, the input *h*_*t*_ and hidden state *z*_*t*_ as well as the transition matrix A and the emission matrix B. (**B**) State occupancy for one feature-channel across 200 trials comprising five blocks. In blocks 2 an 4 the target feature matches that of the channel. For blocks 1, 3 and 5 the target feature is different but can either be in the same dimension (blocks 1,3) or a different one (block 5). Open circles show whether the channel feature was chosen (1) or not (0). The color codes for the inferred probability of being in the Avoid-, Explore-, Preferred- and Persist-state according to the legend. (**C**) For different set shifting conditions, the plots show the inferred probability (y-axis) of the four states from left to right: Avoid, Explore, Preferred, and Persist, to the time at which the target feature goes from a non-match to a match of the channel feature under consideration. The data is averaged across all channels and all subjects. (**D**) Same as C for the different gains/loss conditions.

The example state trace was representative of the overall data. The average state transition dynamics of feature channels successfully recovered the behavioral learning performance (**Fig. 2C, D**). When a block transitioned to a new target feature, the feature channel that corresponded to the rewarded target feature ceased being in an Avoid State, increased transiently to be in an Explore state, and gradually increased the likelihood of being in the Preferred and then the Persist state (**Fig. 2C**). The speed of transitioning from exploring a feature (Explore state) to choosing the feature preferentially over other features (Preferred state) was faster in the New-ID and New-ED condition. This finding is consistent with the behavioral learning curves and with a novelty bias of attending and choosing novel over familar objects (Ghazizadeh *et al*., 2016) (**Fig. 2C**). Additionally, the transitioning of feature channels from Explore-to Preferred- and Persist-states was faster in blocks with the lowest incentives (Gain +2) and highest symbolic punishments (Loss -3) (**Fig. 2D**), which recovers the faster behavioral learning in the Gain +2 / Loss -3 condition (**Fig. 1F**).

### State transitions are coordinated across parallel feature channels

The feature-based learning task presented three objects with six different features in every trial. These six features were either identical (same) to those used in the previous block or they were six novel features not shown before. We therefore extended the HMM model to track the belief states for twelve features in parallel across block transitions. We denote this model the *Parallel Belief States* (PBS) model. The PBS model estimates the state transitions of all twelve feature channels in parallel, using the same four hidden belief states for each channel that were used for the single channel model (*see* above) (*see* Methods, **Suppl. Fig. 1A-C**). Fitting the PBS model to the choices of subjects provides quantitative evidence that during learning subjects explored multiple features in parallel (i.e. multiple feature channels are in an Explore state). Channels of non-target features gradually transitioned to the Avoid state, while the target feature eventually transitions from the Explore to Preferred and Persist states. This state transition dynamic is illustrated across twelve feature channels and several blocks in **Fig. 3A**, showcasing multiple blocks in which some non-target features remain in an Explore state even in trials in which the target feature channels already had moved to a Preferred or Persist state. **Fig. 3B** visualizes the top row of the multiple feature state map in **Fig. 3A** to illustrate the state dynamic of a single feature channel that is initially not the target but becomes the target feature in a later block. This feature channel was in the Explore state in blocks 1, 2, and 5, in the Avoid state in block 4, and quickly transitioned to the Persist state in the sixth block in which it is the rewarded target feature.

**Fig. 3.**
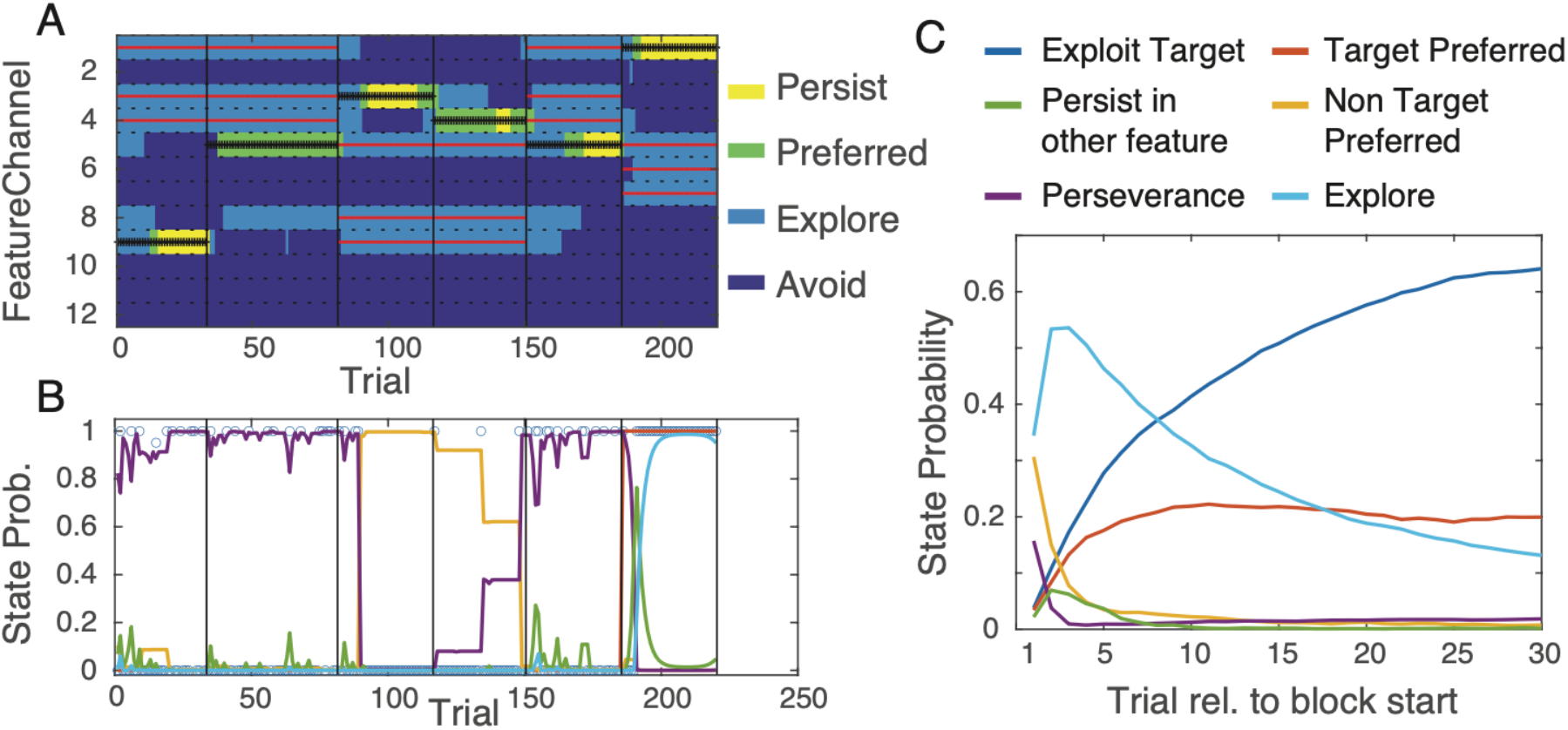
State transitions are coordinated across parallel feature channels. (**A**) Example of the states of 12 feature channels (y-axis, separated by dashed black horizonal lines) over trials of 6 blocks (vertical black lines). The target feature is marked with black plusses. Red lines mark channels with features of a different dimension than the target feature. In block 1, the 9^th^ channel is the target and transitions to the Persist state at a time when the 5^th^ and 8^th^ channel transitioned to Avoid. In block 2 the target channel did not transition to a Persist state. (**B**) State probability of a feature channel that is the target (red line) in the 6^th^ block. Colors indicate probability of being in Avoid (yellow), Explore (purple), Preferred (green), or Persist (blue) states. As target it is quickly chosen and transitions into the Persist state. When the feature is chosen (open circles) in blocks when it is not the target, the channel remains in the Explore state (purple). (**C**) Classifying six separate parallel states from the perspective of the previous and current target channel and average these states across all blocks: Initially the previous target shows some perveration (purple) or remains in a Preferred state (yellow). The new target quickly transitions to an Explore state (light blue) and then from trial 7 onwards into the Exploit state (dark blue).

These state transition dynamics is evidence that learning involves considering multiple features in parallel. According to the PBS model the feature channel states correspond to the degree of feature-based attentional priority. In a task with twelve feature channels and four states per channel this priority map can have up to 4^12^ distinct possible parallel channel states. However, the likelihood of parallel states is constrained by the state dynamics of feature channels that are current targets or were targets in the preceding block. To highlight this constraint we defined six target-based states to discern the state dynamics of the current and previous target feature (**Fig. 3C**). In this schema the current target feature can be in the Persist state (target-based state 1: Exploit Target), in the Preferred state when in some cases other feature(s) are also Preferred (2: Target Preferred), or in a non-preferred state while non-targets are in the Preferred state (3: Non-Target Preferred state). Alternatively, trials can have non-target features in the Persist state (4: Persist in other features), with a special case when the target feature of the previous block remains in the Persist state (5: Perseverance). A final target-based trial state is defined when trials have no feature channel in the Preferred or Persist state but have several features in the Explore state (6: Explore). The occurrences of these target-based states systematically varied with learning (**Fig. 3C**). At the beginning of a block there are few trials with Perseverance (purple) and with preferred choices of a non-target features (yellow). At the same time the Exploration state (cyan) become more likely and then gradually decreases while states that prefer and exploit the target (red and blue) become more prevalent.

The dynamics of the target-based belief states distinguished feature-based learning (**Fig. 4A-D, H**) and gain-loss conditions (**Fig. 4E-G**). Behavioral analysis showed that learning was fastest in the ED-new condition (**Fig. 1E**). The parallel states recovered this ED-new effect, the Exploit Target states curve lying above all other curves (**Fig. 4A**), whereas the Explore state peaks earlier and decays faster (**Fig. 4B**). Similarly, the behavioral analysis showed that the lowest motivational context (+2/-3 condition) led to fastest learning (**Fig. 1F**), which is reflected in parallel channels reaching the exploit target state fastest and with the highest probability (**Fig. 4E**). The parallel state dynamics also show that perseveration is restricted to only a few trials at the beginning of a block when the same features were used in the current block and strongest when the current and past target feature channel were in the same feature dimension (ID-Same condition) (**Fig. 4H**).

**Fig. 4.**
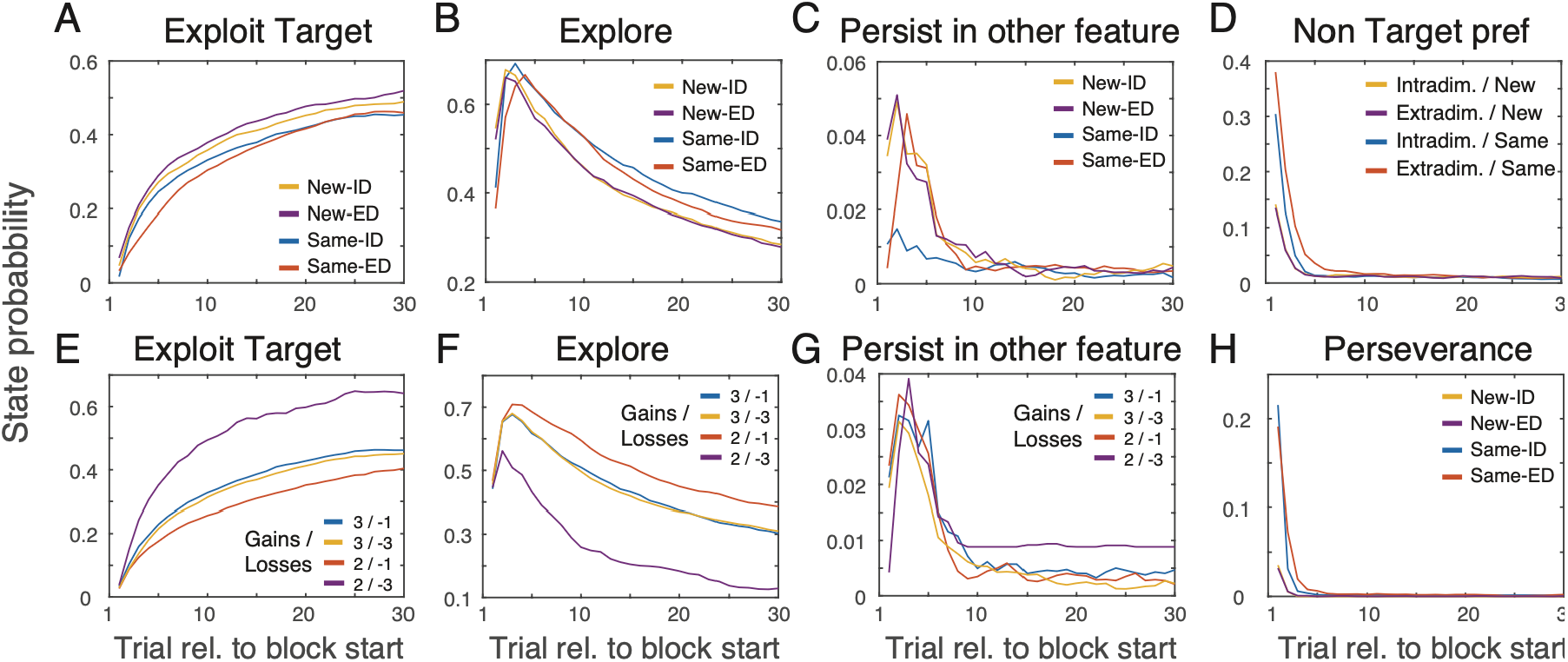
State dynamics of the Parallel Belief State model recovers behavioral learning patterns. (**A-D**) State probability of the target feature channel for Exploit (A), Explore (B) Persist in other feature (C), and Non-target preferred (D) states. Blocks with new features (New-ID, New-ED) show faster rise of Exploit states, while targets using features from the past blocks (Same-ID, Same-ED) have persistently higher Exploration states. (**E-G**) Same format as A-C for the gain/loss conditions. The Exploit state is reached fastest in the +2/-3 condition. (**H**) Perseveration states were rare and more likely in the ‘same’ conditions. State probabilities are averages across all blocks and monkeys.

### Stable feature preferences determine transitioning away from non-target exploration

The preceding results showed how the PBS model captures the state dynamics of flexible learning. We next applied the PBS model to identify stable feature preferences and how they affect learning performance. Previous rule-based set shifting studies have documented that subjects show stable biases of learning faster for certain feature dimensions, preferring e.g. shape over color (Mansouri *et al*., 2020). We found that the PBS model identified stable biases in two ways. First, a subset of features reached the Exploit state earlier and stayed in the Exploit starget state longer than other features, signifying that subjects had a stable bias of learning these features preferably over other features (**Suppl. Fig. 2A-D**). Secondly, these preferred features more likely transitioned away from the Avoid state to the Explore, Preferred and Exploit states during learning than the least preferred features (**Suppl. Fig. 2D-G**). These findings show that the PBS model recovers stable biases in learning, suggesting that subjectively preferred features are faster and more likely explored and exploited (**Suppl. Results**).

## Discussion

We showed that the learning of the relevance of visual features is accounted for with a parallel belief state (PBS) model that allows each visual object feature to be in one of four discrete belief states. The four states are described by the probability of choosing the feature, 1: near 0 percent for ‘Avoid’; 2: 33% for ‘Explore; 3: between 40 and 80% for ‘Preferred’; 4: close to 100 % for ‘Persist’. During learning multiple features are in an Explore state, while successful learning is completed when the belief state of the target feature channel is the Persist state while non-target features are avoided (Avoid state). The PBS model recovered effects of selection history and motivation on learning. The Persist state was reached slower for target features that were non-target features in the previous block (i.e. Same ED or Same-ID), which reflects a selection history effect (Awh *et al*., 2012; Anderson *et al*., 2021). In contrast, Persist states were reached faster when subjects anticipate higher costs (loss) of making errors, which reflects a motivational incentive to avoid punishment and is consistent with previous findings (Boroujeni *et al*., 2022a; Boroujeni *et al*., 2022b).

### Parallel Belief States represent a feature-based attentional priority map

One appeal of a Parallel Belief State model to account for learning behavior is the rich information carried by each state. The belief state of a feature is at the same time a reflection of the history of choices and outcomes experienced with that feature and a forward looking probability that the feature will be considered in future choices. According to this formulation the belief states resemble action-value states in reinforcement learning (RL) frameworks. In RL models action-value states are estimated using prediction errors in order to predict rewards and (potentially discounted) values of future rewards of actions in order to maximize future rewards (Sutton & Barto, 2018). Across different actions subjects can take the action state landscape thus represents the behavioral policy an agent follows. In analogy to the action state landscape reflecting a behavioral policy, the landscape of feature belief states in the PBS model can be considered attentional priorities of object features. This feature-based attentional priority constitutes a policy that determines whether features will be avoided, explored, preferred, or exploited. This rank ordering of states corresponds to the degree of attentional priority. Subjects are expected to attend most (least) likely features that are in the Exploit (Avoid) state while features in the Explore state have intermediate attentional priority and are occasionally considered.

Beyond the similarity of action states in RL models and the feature belief states described here, there are also major differences. First, the PBS model is an observational model that does not specify underlying mechanisms of the policy updates but rather aims for a statistically optimal prediction of feature choices. While RL models propose roles of reward- and state-prediction errors for the state update, the PBS model uses a transmission model that explains transitions of feature states by a combination of previous choices, outcomes, and current hidden states. In our specific instantiation of the PBS model the behavioral data were best fitted by considering the current state (obtained by previous choice and outcome in conjunction with its own past states), but without the need to consider past reward outcomes in the emission model for predicting the choice. This finding shows that the ongoing hidden belief states represented the history of recent outcomes without the need to consider outcomes explicitly for the choice.

A second, related difference of the RL and PBS approach is that the PBS model optimizes the states to best reproduce (predict) the feature choices of subjects, while RL frameworks aim to maximize rewards. By focusing on fitting choices of subjects, the PBS model can transition states abruptly when the pattern of choices change, while value learning in the RL framework is generally more gradual with continuously assigning a higher value to a state (i.e. feature value) depending on a learning rate parameter (Soto *et al*., 2023).

### Parallel Belief States may be enacted in neuronal feature-state encoding ensembles

Recent studies of neuronal representations of attentional top-down informations that neuronal populations encode relevant target features in lower dimensional spatio-temporal activity in fronto-parietal and fronto-striatal neuronal populations (Johnston & Fusi, 2023; Jahn *et al*., 2024; Tian *et al*., 2024). The Parallel Belief State model is consistent with these findings. We hypothesize that feature-specific states are represented in a recurrent network of neurons that have a unique firing pattern for each feature-state pair (**Fig. 5A**). When a target feature has been learned its Preferred- or Persist-state is assumed to be represented in a unique firing pattern. When the feature stops being the target and ceases to be chosen and rewarded, it transitions to Explore and Avoid states. State transitions are effectuated by a weakening of connectivity of neurons forming the previous state and a strengthening of connections representing the new state (**Fig. 5A**). This neuronal implementation provides explicit hypotheses about why novel object features are learned faster and how motivational contexts can speed up state transitions and can be tested using population analysis methods (Kobak *et al*., 2016; Williams *et al*., 2018; Williamson *et al*., 2019). When a novel feature is presented it is expected to cause a higher firing rate increase than a feature that was previously shown and is in an Avoid state, i.e. a previous distractor feature (**Fig. 5A**, blue colored feature-state). The higher initial response accelerates the state transitioning of the neuronal ensemble representing feature-states for those features, which would result in faster behavioral state transitions for new rather than previously experienced feature states and accounts for the faster learning in New-ID/New-ED than in the Same-ID/Same-ED conditions.

**Fig. 5.**
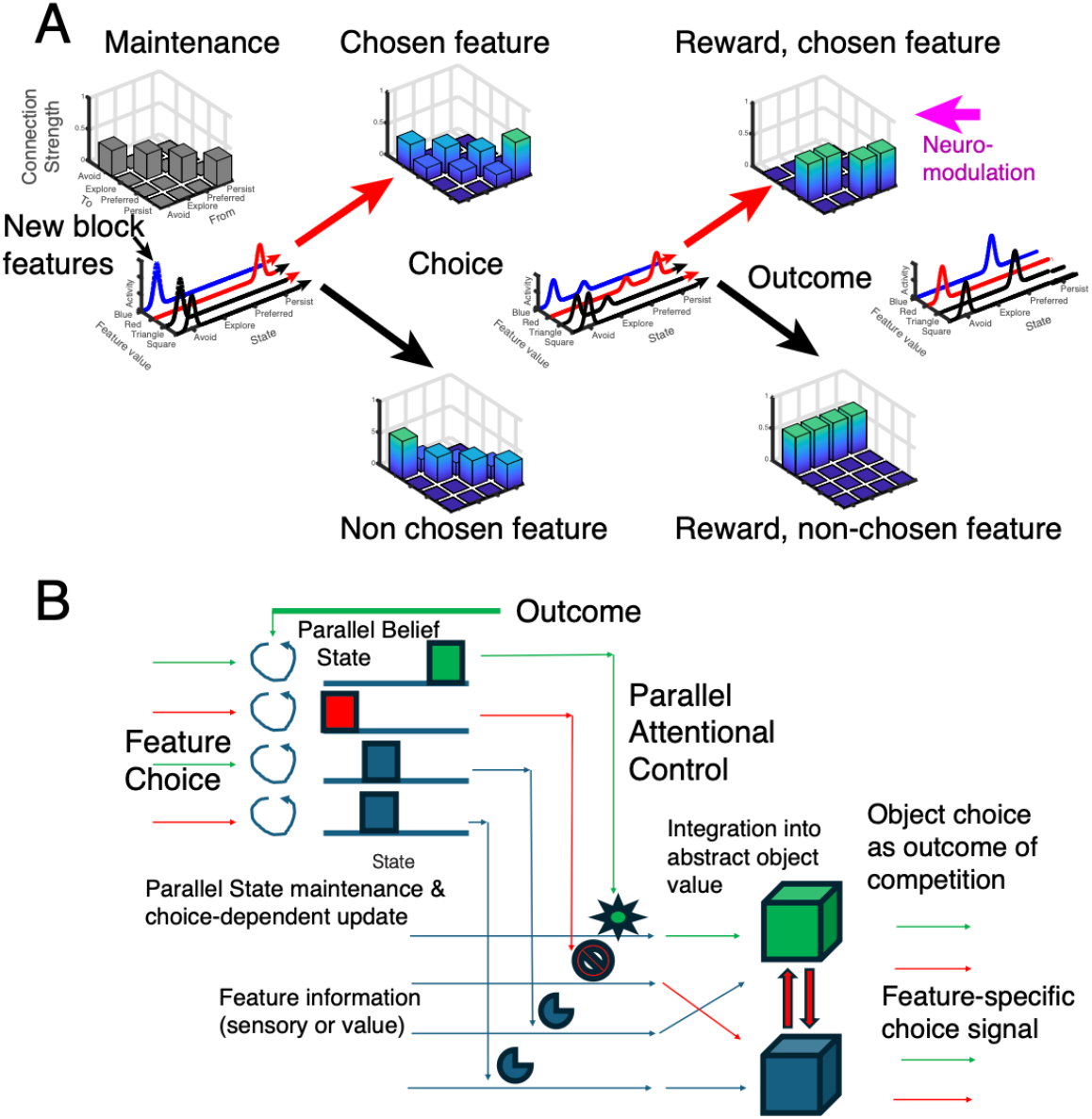
Updating the belief state of parallel feature channels and the role of attentional control to integrate feature-specific belief states into an abstract value for each object. (**A**) We hypothesize that *Parallel Belief States* are represented in populations of neurons labeled by the feature (*y-axis*) and state (*x-axis*). Prior to a choice (*left*), belief states for all available features are reflected in firing rates (*bell-shaped curve*). Activity is higher for features that are new (*blue*) which is a putative mechanism for faster updates. The neuronal population encoding feature-states is interconnected, represented as a 3D bar graph, with efferent (*x-axis*) and afferent (*y-axis*) connections and activity bumps reflecting attractor states. State transitions are initiated by a choice (*middle column*) by spreading activity from the current to the neighboring state, e.g. away from the Avoid state for features that were chosen (*top matrix*), and towards Avoid state when not chosen (*bottom matrix*). The outcome (*right column*) triggers the final state update: Chosen and rewarded features transition to a Preferred state, the non-chosen features transition to the Avoid state. Motivational factors like loss avesion may act on neuromodulators that change the speed of state transitionining through changes of the connection matrix. (**B**) Hypothesized integration of Parallel Belief State channels into an abstract value for each object. When a channel state is Persist (*green star*), its feature information is selected to pass, while channels in the Avoid state (*red stop sign*) are blocked and gated-out. Feature channels that are in the Explore state, are integrated but with a reduced weight (*blue pacman symbols*). A choice is made through competition (reciprocal red arrows) among belief state channels, favouring objects with more features in the Persist state. The choice tags feature channels (*green, bottom right*) and feeds them back into the parallel belief state network (top left).

We hypothesize that motivation influences the speed of learning with a different mechanism that speeds up plasticity changes within the neuronal population encoding feature specific belief states. Subjects transitioned faster from Avoid or Explore to Preferred and Persist states when anticipating higher loss for erroneous choices (**Fig. 2D, 4E,F**). Faster state changes could be realized by neuromodulators that attenuate connection weights of neurons coding for the previous feature state, and enhancing connection weights among neurons encoding the novel feature belief state (Frémaux & Gerstner, 2016).

The outlined neuronal dynamics of state transitions of the PBS model also predicts a discrete sequence sof state changes with each behavioral choice (**Fig. 5A**). Attentional priorities reflect the initial state that guides a choice, while during the choice itself the neuronal representation changes so that neuronal ensembles coding for the states of the chosen features already activate to impose a state transitioning prior to receiving feedback about gains and losses. When the outcome is revealed the recurrent connectivity matrix updates those states that map onto the chosen and rewarded feature states. According to this scenario, parallel belief states reflect an attentional priority prior to a choice and undergoes state updates during the choice and outcome periods to arrive at new parallel belief state changes after each choice-coutcome event.

One important question for future work is how hidden belief states of invidual features affect choices. We hypothesize that the individual feature-specific belief states are integrated into an abstract object value prior to influencing a behavioral choice. **Fig. 5B** outlines that this integration is well conceived of as an attentional control process that gates out feature state channels that are in an Avoid state and selects and amplifies channels that are in an Explore, Preferred and Persist state. This gating process shares similarity to the softmax selection used in reinforcement learning models that aggregate values of features of an object and chooses the objects with higher aggregate values (Womelsdorf *et al*., 2022).

## Conclusion

We proposed a model of discrete belief states that quantifies how attentional priorities are distributed across multiple features during learning. The strength of this observational model is a concise delineation of states and their transitioning during a complex feature-based learning task. This landscape of states and their updating constrains mechanistic hypothesis on how feature-specific beliefs are represented (Kemp & Tenenbaum, 2008; Ostojic & Fusi, 2024), how states are maintained in working memory (Baddeley, 2012), how multiple feature states are integrated, selected and gated during adaptive behavior (Miller & Buschman, 2013), and how separable states compete for choices (Shadlen & Kiani, 2013; Hanks & Summerfield, 2017). These cognitive processes require computational motifs that map onto neuronal circuit structure, such as, for instance, a winner-take-all motif with inhibitory neurons (Usher & Niebur, 1996; Rutishauser *et al*., 2012). The strength of the PBS model is that enables concise testing of computational motifs of the major cognitive control processes underlying adaptive behaviors.

## Methods

### Experimental Procedures

All animal and experimental procedures were in accordance with the National Institutes of Health Guide for the Care and Use of Laboratory Animals and approved by the Vanderbilt University Institutional Animal Care and Use Committee (M1700198-01).

Four pair-housed, adult male macaque monkeys performed the experiments using touchscreens in their home cage. The 19.5’’ touchscreens were mounted in kiosk testing stations that contained a fluid reward pump and a sipper tube for delivering fluid reward during task performance (Womelsdorf *et al*., 2021). On each weekday, monkeys were individually given access for 90-120 minutes to an apartment cage with the mounted Kiosk Testing station and could chose to engage with the touchscreen that controlled the cognitive task. Stimuli, reward, and timing of the task was controlled by M-USE (Multi-task Unified suite for experiments), which is a unity-based platform for psychophysical experiments (Watson *et al*., 2023). Monkeys were fluid-controlled and earned water reward by performing the task.

### Task paradigm

The task required monkeys to choose on every trial one of three objects to earn reward-tokens that were regularly cashed out for fluid rewards. On each trial only one of the objects contained a visual feature that was associated with reward (Fig. **2A**). Monkeys had to learn which object feature is rewarded through trial-and-error. When they made a choice, a visual halo appeared behind the chosen object in yellow if the choice was correct, and in blue otherwise. Correct choices were followed by green circles (tokens) shown above the chosen object. Tokens moved up towards a token bar and populated previously empty token-slots (Fig. **2A**). After an incorrect choice and blue halo, grey tokens appeared above the chosen object, moved upwards to the token bar and the grey tokens were subtracted from the already existing green tokens at the token bar. The token bar started at the beginning of the experiment and in each new block of trials (see below) with three already available tokens and 5 empty token slots. When monkeys collected eight tokens the token bar flashed, the monkey received water reward, and the token bar was reset to contain three default tokens.

### Block transition types

After blocks of 32-65 trials with the same rewarded target feature, the rewarded target feature changed. The experiment pseudo-randomly determined for the new block whether the newly reward target feature was (1) a feature of the same visual feature dimension as the previous target such as a switch from a red to a green color, or from an oblong to a pyramidal shaped object (intra-dimensional (ID) block transitions), or (2) a feature from a different visual feature dimension as the previous target feature, such as a switch from a red color to an oblong shape (extra-dimensional (ED) block transition). In addition to the ID / ED transitions a new block could involve objects (3) with the same visual features (Familiar Objects: same) or (4) with a new set of visual features (Novel Objects: new). In each block the three objects differed along two visual feature dimensions (e.g. different colors and shapes). The objects were so-called Quaddle objects that allowed to parametrically vary the four feature dimensions color, shape, surface patterns, and arm shape (Watson *et al*., 2019).

The experiment also varied the motivational context between blocks. Within a block, monkeys either earned 2 or 3 tokens for a correct choice, and they either lost 1 or 3 tokens for an incorrect choice, which combined to four distinct contexts (Gain 3/Loss 1; Gain 3/Loss 3; Gain 2/Loss 1; Gain 2/Loss 3). The motivational contexts remained constant within a block and varied pseudo randomly within an experimental session so that each context was equally often assigned to each of the four block transition types (same/new and ID/ED transition types).

### Learning state analysis

The standard learning curves are determined by averaging the outcome (reward no=0, reward yes=1) across blocks, with the trial index aligned on the block start. An alternative method, which does not require averaging, considers that the outcome is generated by a Bernoulli process with a probability due to chance (no learning) plus a learning state ‘x’. The learning state for each trial can be inferred from the outcome sequence using the EM algorithm (Dempster *et al*., 1977), and can be smoothed by using the entire sequence to estimate the state of a trial t, rather than only the outcomes preceding the trial. The learning state is achieved when the performance is above chance, that is the learning state is nonzero, with a probability exceeding 0.95. We used the MATLAB implementation given in (Smith *et al*., 2004) to infer the learning state. For each block we determine whether a learning state is achieved, and if so, at which trial it was achieved (Fig. **1 E,F**).

To determine whether the experimental conditions (ED/ID, Same/New, Gain and Loss coditions) affected learning behavior we calculated for each condition the learning curve averaged first across all blocks irrespective of learning state, for each subject, and averaged these over all subjects.

### Hidden Markov Model (HMM)

During the experiment three objects are presented, with two feature dimensions (e.g. color, shape), for each of which there are three feature values that are exclusively distributed across the objects. This means during a block there are 9 possible objects, 3 of which of have the target feature, that can be presented in 36 unique combinations of 3 objects. Each trial is therefore typically described in terms of what was the target feature, which three objects were presented, which one was chosen and what was the outcome (reward or not; simplified from the token gain or loss actually used). We fit these experimental results using an input-dependent hidden Markov process – here referred to as GLM-HMM because the emission matrix can be viewed as a generalized linear model (Ashwood *et al*., 2022). The model analyzes each feature value independently, each trial is described by whether the feature was present in the chosen object (‘feature chosen’, *c*_*t*_ = 1) or not (‘feature not chosen’, *c*_*t*_ = 2) and whether the chosen object was rewarded (*r*_*t*_ = 1) or not (*r*_*t*_ = 0). The goal of the model is to predict the choice (*c*_*t*_) on trial t based on a hidden state (*z*_*t*_) and the input, which is coded as follows *h*_*t*_ = *c*_*t−*1_ + 2(1 *− r*_*t−*1_) in terms of a number between 1 and 4. This numerical input can also be represented symbolically as 1:C+R+, 2:C-R+, 3:C+R-, 4:C-R-. The hidden state can be interpreted as a belief about whether the feature modeled is the target feature, but its meaning is more properly determined by the emission matrix introduced below.

We can write down the probability for the entire set of observations (*c*_1*:T*_) across trial 1 to T, and hidden states (*z*_1*:T*_) given the inputs (*h*_1*:T*_) and a particular model m with parameters *θ*_*m*_.

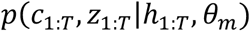

We need to optimize this, but we do not know the hidden state. Expectation-maximization uses a trick to eliminate the dependence on the hidden states, it assumes that an estimate for the parameters, *θ*_*old*_ (for clarity we suppress the dependence on the model), is available, hence, the probability distribution for the hidden states can be calculated:

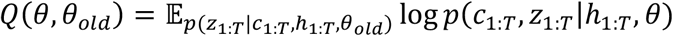

The Q is the estimate for the log-likelihood of the observations. The first step is to perform the average (E step) and then maximize the Q (M step) as a function of *θ*.

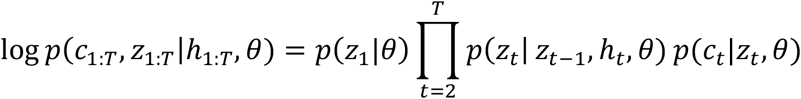

The first term is the prior on *z*_1_, which is a vector with a probability for each possible state, *z*_1_ = 1 to *z*_1_ = *n*_*s*_, denoted by *π*. The second term describes the transition from the previous hidden state to the current hidden state, which is a *n*_*s*_ × *n*_*s*_ matrix that can be dependent on *h*_*t*_, which means there are *n*_*i*_ = 4 of them, which we denote by *A*^1^ to 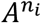. The third term is the emission model, which provides the probability for a choice given the current state, and this may also dependent on the input. The emission model is therefore a set of *n*_*i*_ 2 × *n*_*s*_ matrices, denoted by *B*^1^ to 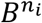. Taken together the parameters are 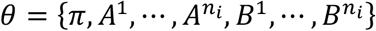. The sum of the elements of *π*, and each column of A and B has to add up to 1. This means we have in total (*n*_*s*_ *−* 1) + *n*_*i*_*n*_*s*_(*n*_*s*_ *−* 1) + *n*_*i*_*n*_*s*_ parameters. We evaluate four types of models:

- No hidden states, in which case, only the emission matrix (with *n*_*s*_ = 1) is a parameter, i.e. *n*_*i*_ parameters. Here we also vary how much back in trials the input looks.
- No input dependence of the emission matrix: (*n*_*s*_ *−* 1) + *n*_*i*_*n*_*s*_(*n*_*s*_ *−* 1) + *n*_*s*_ parameters
- No input dependence of the transition matrix: (*n*_*s*_ *−* 1) + *n*_*s*_(*n*_*s*_ *−* 1) + *n*_*i*_*n*_*s*_ parameters
- Full model: (*n*_*s*_ *−* 1) + *n*_*i*_*n*_*s*_(*n*_*s*_ *−* 1) + *n*_*i*_*n*_*s*_ parameters

We now describe the algorithm used for the E step. It utilizes the following quantities:

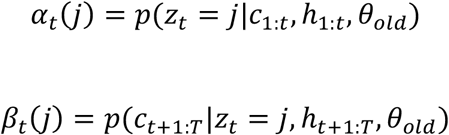

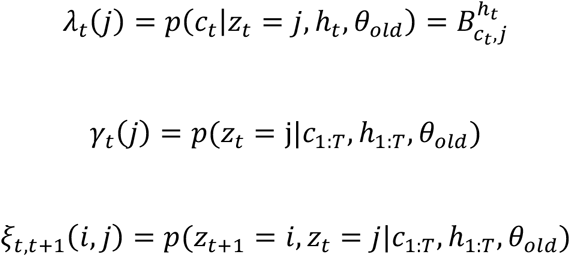

For *α* a forward sweep is used, starting at *α*_1_ with the prior *π*,

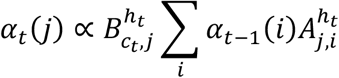

The proportionality sign indicates that this quantity needs to be normalized, so that the sum across j is one.

For *β* a backwards sweep is used, starting with *β*_*T*_ equal to unity,

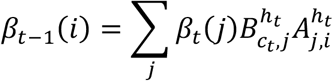

Then we have

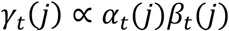

And

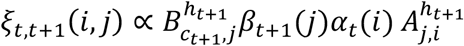

In these expressions the A,B and *π* are the old parameters. With these quanties we can now write down Q:

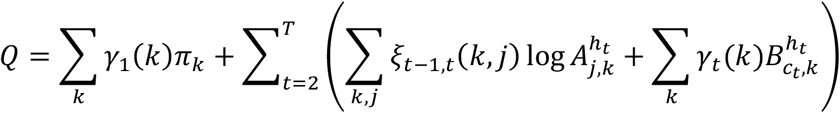

This yields the M update, by setting its derivative to zero while applying Lagrange multipliers to enforce the normalization of the columns of the matrices:

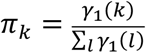

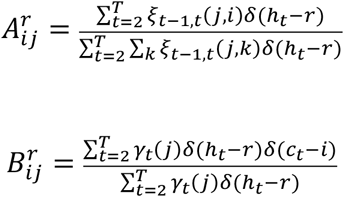

This sequence of steps is repeated until Q does not decrease appreciably anymore, in our hands we found 300 steps to be adequate.

### Preprocessing the dataset for modeling

Each block is characterized by a target feature, which is always present in one of the three presented objects, the reward structure in terms of how many tokens are gained (lost) with a correct (incorrect) choice and a blocktype regarding how the target feature and presented objects changed relative to the previous block. For each channel feature f, we translate the chosen object and outcome trial sequence into *c*_*t*_ = 1 when the chosen object contained the feature, and *c*_*t*_ = 2 when it did not, the previous choice *c*_*t−*1_ and previous outcome *r*_*t−*1_ is translated in an input on the current trial *h*_*t*_ = *c*_*t−*1_ + 2(1 *− r*_*t−*1_). In order to learn all the states properly, we found that it is useful to make sure that all states occur often enough. The ‘Persist’ state occurs when the channel feature matches the target feature. Hence, we found all blocks for which this was the case, for each target block we prepended the previous block and then stacked the resulting two blocks behind each other to obtain the entire trial sequence. We did this for each channel feature for which there were more than 2000 trials available and fitted the combined data set, containing more than 100 thousand trials using the EM procedure.

We were also interested in following the state transitions in multiple parallel channels simultaneously, for which the aforementioned procedure is inappropriate. In a given block there are six features present in the objects, when in a subsequent block a new dimension is introduced (in the condition ‘new’), there are three or six new features added. We want to track these block transitions, but avoid having to track the states of too many features simultaneously because then we will not get appropriate statistics on each state, hence we cut the trials into groups of consecutive blocks in which no more than 12 unique features are present. Each group undergoes the forwards-backwards sweep independently (E step), but their statistics are combined into a single dataset for performing the M step.

In a seperate fit, for the independent channel analysis, we give each block an additional label, and assign a separate set of transition matrices to each label. The label can reflect the block type, or the feature in terms of its long-term bias, so that we can assess the effects of these conditions on the transition matrices.

### Preprocessing parallel feature channels

The Parallel Belief State (PBS) model infers the state of multiple feature-channels simultaneously. To achieve this we aggregated the blocks by selecting sets of consecutive blocks (epochs) where at most 12 distinct features are presented, rather than the total number of 37 features across the entire data set, to make sure that each channel cycles through all available states during an epoch. In addition, we fixed the emission matrix B to enforce a uniform definition of states across subjects.

### Definition and analysis of states, emission and transmission matrices during learning

We use the Hidden Markov Model (HMM) to describe learning the rewarded feature in terms of the state of feature-channels, for each feature separately. The HMM infers the states and determines the state transition during the current trial as a function of an input which represents the choice and outcome on the preceding trial, a GLM-HMM (Bengio & Frasconi, 1994). The mechanics of the model is summarized in **Figure 2A**. The model accounts for whether a feature was present or not in the chosen object by maximizing the probability of the observed choices {*c*_1_, ⋯, *c*_*T*_} for a given channel feature given the history of past choices and the resulting outcomes. Specifically, the choice *c*_*t*_ (value 1 if the feature is chosen, 2 if it is not chosen) on trial t is predicted based on the current input *h*_*t*_ = *c*_*t−*1_ + 2(1 *− r*_*t−*1_) which reflects the choice and outcome on the preceding trial encoded as a number between 1 and 4. For example, if the feature was chosen (*c*_*t−*1_ = 1) and the outcome was a reward (*r*_*t−*1_ = 1), then *h*_*t*_ = 1. This input directly influences the predicted choice via the input-dependent emission matrix 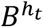, which links the current hidden state *z*_*t*_ which reflects the longer-term history to a probability of the choice *c*_*t*_ via the matrix element 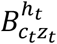 of this matrix. The input dependent transition matrix 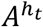, updates the hidden state to reflect the most recent available choice/outcome, which reflects the learning dynamics of the subject. This new hidden state is not directly accessible for the observer, instead it must be inferred, which yields a probability *γ*_*t*_(*j*) for *z*_*t*_ to be in state j when taking into account both the past and the future.

The goodness of the model, i.e. of the prediction, is assessed by the likelihood of the actual choice under the model and is maximized via an EM procedure. We used the log likelihood (LL) of the entire sequence of choices, normalized by the number of choices, its value should exceed that of chance 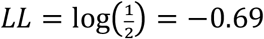 (note that we use a natural e-based log). The LL increases with iterations. The task used 37 different features across all experimental sessions, which allowed constructing a dataset out of 37 feature choice-outcome sequences. In order to assess variability we created 50 new data sets by choosing 37 features randomly with replacement. For each of these data sets we ran the procedure 10 times for different initial values for the parameters (the matrices A, B as well as the prior on the hidden states, *π*), to avoid the EM procedure getting stuck in local maxima. The resulting 500 curves are shown in (**Suppl. Fig S1A**). The model converged typically in about 100 iterations, we therefore performed a minimum of 300 iterations for our analysis. For each data set we choose the initial condition with the largest LL, sorted the resulting 50 runs and color coded all corresponding curves according to the rank, from dark blue for low to yellow for the highest rank (**Suppl. Fig S1A**). The variability due to the initial conditions is smaller than that due to the specific combinations of features in the data set.

We set out to find the best set of parameters for the model, in terms of the dimensionality of the hidden state, *n*_*s*_, the depth of the history *h*_*t*_ (how many previous trials are incorporated, in our standard formulation this was 1) and the best model structure in terms of whether both A and B are input-dependent. We explored *n*_*s*_ = 2, 3, 4 and 5 and found four states to be optimal, as considering 5 states only led to a marginally better fit, but it had many more parameters (data not shown). We then ran fits with *n*_*s*_ = 4 and (1) the full model (label “full”), (2) with only the A matrix input dependent (“noB”) and (3) only the B matrix input dependent (“noA”) (**Suppl. Fig S1B**). Both the full and noB model variants were similar in LL values but both were better than noA. The full model had a slightly better BIC than the noB model, but this significance was significant only with a t test but not with rank sum test. We therefore focused on the noB model in the main text.

The model well fit to each of the four subjects, with subject M1 yielding highest LL (**Suppl. Fig S1C**). This corresponded to subject M1 showing the highest fraction of blocks in which a learning state was acquired, i.e. with the most trials where choice probabilities were high, which gives rise to a higher LL (which is the log of the choice probability).

The top row 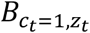 of the B matrix obtained in the noB fit, shows the probability of choosing the feature for each state. We define the states as “Avoid”: do not choose the feature (lowest probability the feature is chosen); “Explore”: choose the feature randomly, with about 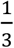 probability; “Preferred”: choose the feature more often, typically between 0.4 and 0.8 probability; “Persist”: (almost) always choose the feature (highest probability the feature is chosen). The state has therefore a clear interpretation in terms of predicted feature choice. We also consider the B matrices obtained in the full fit in two different ways. First, we indicate the effect of input which shows that in the Avoid state, the feature will still be chosen when it was chosen on the previous trial (C+) and when that choice resulted in a reward (R+) (**Suppl. Fig. 1D**). Likewise, in the “Persist” state, a feature will also be chosen when it was chosen in previous trial most likely when it was also rewarded (C+R+), but is still chosen with half the probability when it was chosen and not rewarded (C+R-). Second, we can order them according to input and color code the state (**Suppl. Fig. 1E**). This immediately shows for which inputs, the state has an overrriding influence, for instance, for the not-chosen, not-rewarded (C-R-) input, the state determines the choice.

We evaluated the role of the A matrices on the probability of reaching all other states j from the Persist state 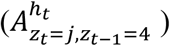 and those that are reached from the Avoid state for two inputs, when the feature was chosen on the previous trial and either was rewarded (C+R+) or not (C+R) (**Suppl. Fig. 1E**). For a rewarded choice of a feature (input C+R+), the model stays in the Persist state, whereas it moves out of the Avoid state. Overall, the model moves from Avoid most often to the Explore and otherwise Persist state, avoiding the Preferred state. On the other hand, after a non-rewarded choice (C+R-), you expect a move out of the Persist state, whereas you expect to stay in the Avoid state. Consistent with this expectation, the model most frequently moves from the Persist to the Explore state, bypassing the Preferred state. The form of the A matrices for these conditions were remarkably similar across subjects (**Suppl. Fig. 1F,G**).

Taken together, we use the model with a A matrix (noB model) for the behavioral analysis of the behavior (see main text). The input dependent A matrix encodes the trial history into the hidden state at a slower time scale (gradually) than the B matrix, which has an immediate impact on feature choice. If a model would include both, A and B, both matrices would influence choices and cause competition between causes, and hence, more variability across fits and a more difficult interpretation. We focused on the noB model also to avoid this competition, which allows for a more straightforward interpretation of the states and their updates via the A matrices.

## Data and code availability

Data and custom programming code for analysis will be made available on Zenodo upon publication.

## Acknowledgments

We thank Adam Neumann for invaluable help training and testing the nonhuman primates. This work was supported by the National Institute of Mental Health (R01MH123687). The funders had no role in study design, data collection and analysis, the decision to publish, or the preparation of this manuscript.

## Supporting Information for

### Supplementary Results

#### Inferring stable preferences of features during learning

To identify whether subjects had preferences for specific features that may have biased learning speed we compared how often each feature was chosen, that is, how often it was present in the chosen object, normalized by the number of trials on which the feature was presented in one of the blocks (**Suppl. Fig. S2A**, solid lines). As this measure could be influenced by differences between features in how many trials a feature was the target, we recalculated the fraction by considering only the trials in which the feature was not the target (**Suppl. Fig. S2A**, dashed lines). For both calculations, and across all subjects, there was a difference in the fraction of trials on which the presented feature was chosen. We used the second method (excluding trials where it was the target) to order and color code each feature from preferred (blue) to least preferred (yellow) (**Suppl. Fig. S2B**). The preferredness had an effect in how quickly the Persist state (Exploit Target state) was reached after the target changed at the beginning of a block, and when the accuracy plateaued towards the end of the block (**Suppl. Fig. S2B**, example data for subject M1).

We grouped the ten most preferred features (MPF) and the seven least preferred (LPF) and calculated the mean Persist-state curves (**Suppl. Fig. S2C**), which showed that there was a similar difference between MPF and LPF Persist-state curves across subjects. The choice for 10 in one group and 7 in the other was based on knee in the chosen fraction curves for M1 (**Suppl. Fig. S2B**) and was kept the same across all subjects.

There are two alternative ways to evaluate the effect of preferredness on learning dynamics. One is to fit the blocks across all features with a single transition matrix and infer the states, from which differences in state dynamics across features emerge, as shown in **Suppl. Fig. S2B** and **C**. An alternative is to fit a different transition matrix for each group and compare the matrices to assess for differences in learning dynamics. We selected a subset of matrix elements for further analysis, specifically we considered the states reached from either the Avoid or the Persist state, when on the previous trial the feature was chosen (C+) and either rewarded (R+) or not (R-). This tells us how positive and negative information is used to change the state, that is, the confidence in whether the feature is the target or not.

When a feature was chosen and the object choice was rewarded (C+,R+), you would expect the state to remain in Persist (**Suppl. Fig S2D**) and move away from Avoid (**Suppl. Fig S2E**). This is indeed what happens, except that for MPF there is some small leakage away from Persist. There are more differences between MPF and LPF regarding where to the state switches from Avoid. For MPF the bulk is to the Explore state for M1, M2, and M4, whereas for M3 it is mostly to Preferred, and for M2 also to Persist. In contrast, for LPF it is more spread out across Random and Preferred states, even reaching Persist.

When a feature was chosen and this choice was not rewarded, you would expect transitions away from Persist and the state to stay at Avoid. The bulk of MPF moves from Persist to Random (M1,M2 and M4) or Preferred (M3), whereas LPF also does that, but even has some transitions towards Avoid. Hence, negative outcomes drive LPF faster to Avoid than MPF. Interestingly, LPF mostly stay at Avoid, but MPF transition to Random as well. Perhaps this reflects optimism that the MPF could still become the target despite the negative outcome on the previous trial.

Taken together, these two analyses show that feature preferredness affects learning dynamics, with both negative and positive outcomes affecting stronger state changes for LPF, but nevertheless resulting in a slower learning when considering the Persist state curves.

## Supplementary Figures

**Suppl. Fig S1.**
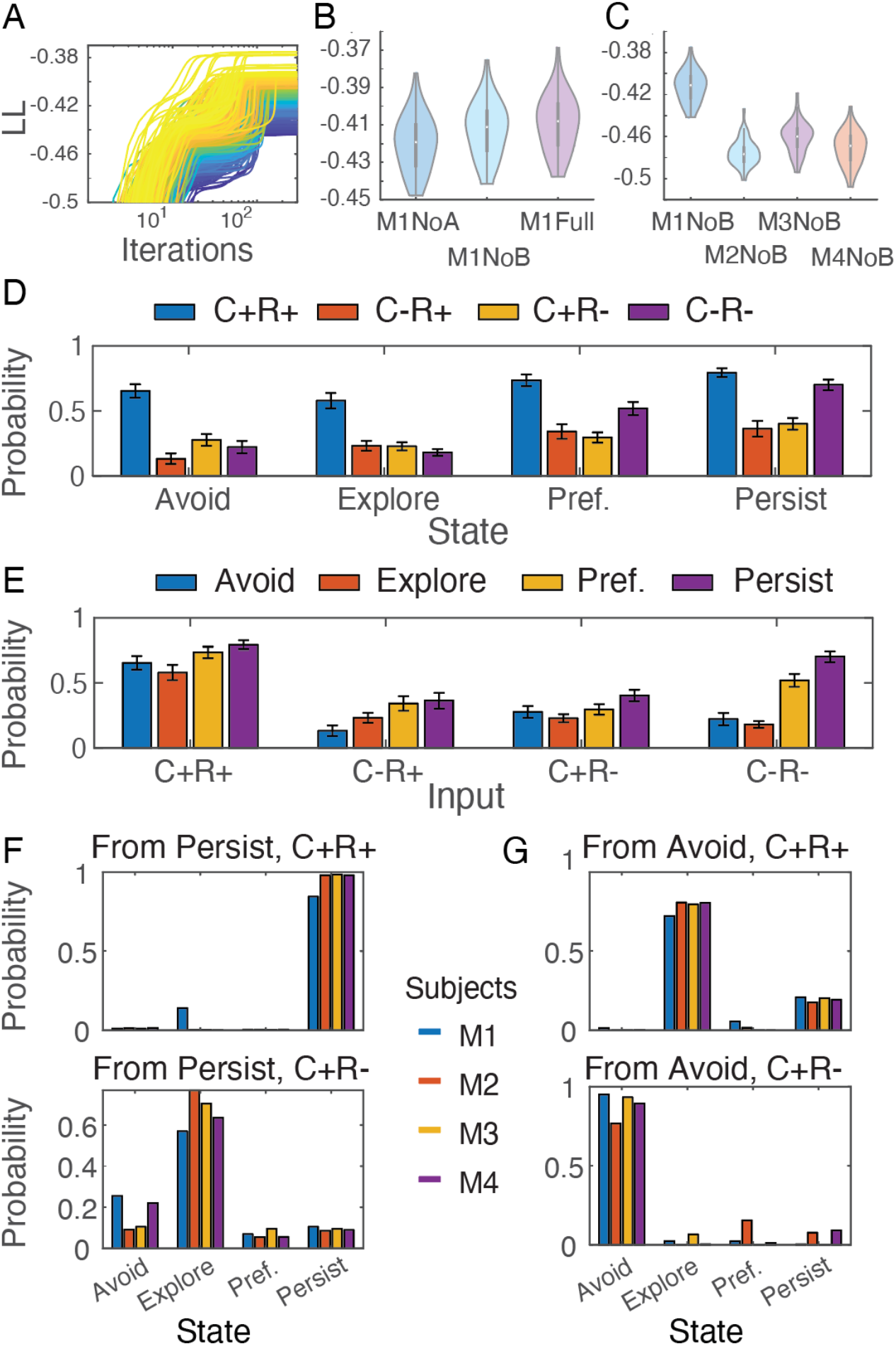
The best model for predicting feature choices had an input-dependent transition matrix A and an input-independent emission matrix B. (**A**) The log-likelihood (LL) of the observed sequence of feature choices given the current parameter settings as a function of iteration. We show 50 different bootstrapped data sets, each started from 10 different initial positions. The curves are color-coded according to the rank of the LL of the converged best (across initial conditions) model. Higher, less negative values (more yellow), correspond to better solutions. (**B**) LL across 50 data sets, for three model structures, “full”, with both A and B input dependent; “noA”, with only B input dependent; “noB”, with only A input dependent; with a fixed number of states, *n*_*s*_ = 4. The “noA” condition is worse, whereas the noB and full conditions are comparable. (**C**) All subjects could be fit, with M1 yielding significantly better fits than the other 3 subjects. The violin plots for *n*_*s*_ = 4 hidden states and the “noB” model structure are shown. (**D,E**) The emission matrix B for the full condition ordered to either highlight the effect of input given a state (*E*) or the effect of state given an input (*F*). As the columns of the 2 × *n*_*s*_ B matrix are normalized, we only plot the upper row. Example data for M1, errorbar is across bootstraps. (**F,G**) The transition out of (*F*) Persist and (*G*) Avoid state when the input was (top) C+R+ or (bottom) C+R-.

**Suppl. Fig S2.**
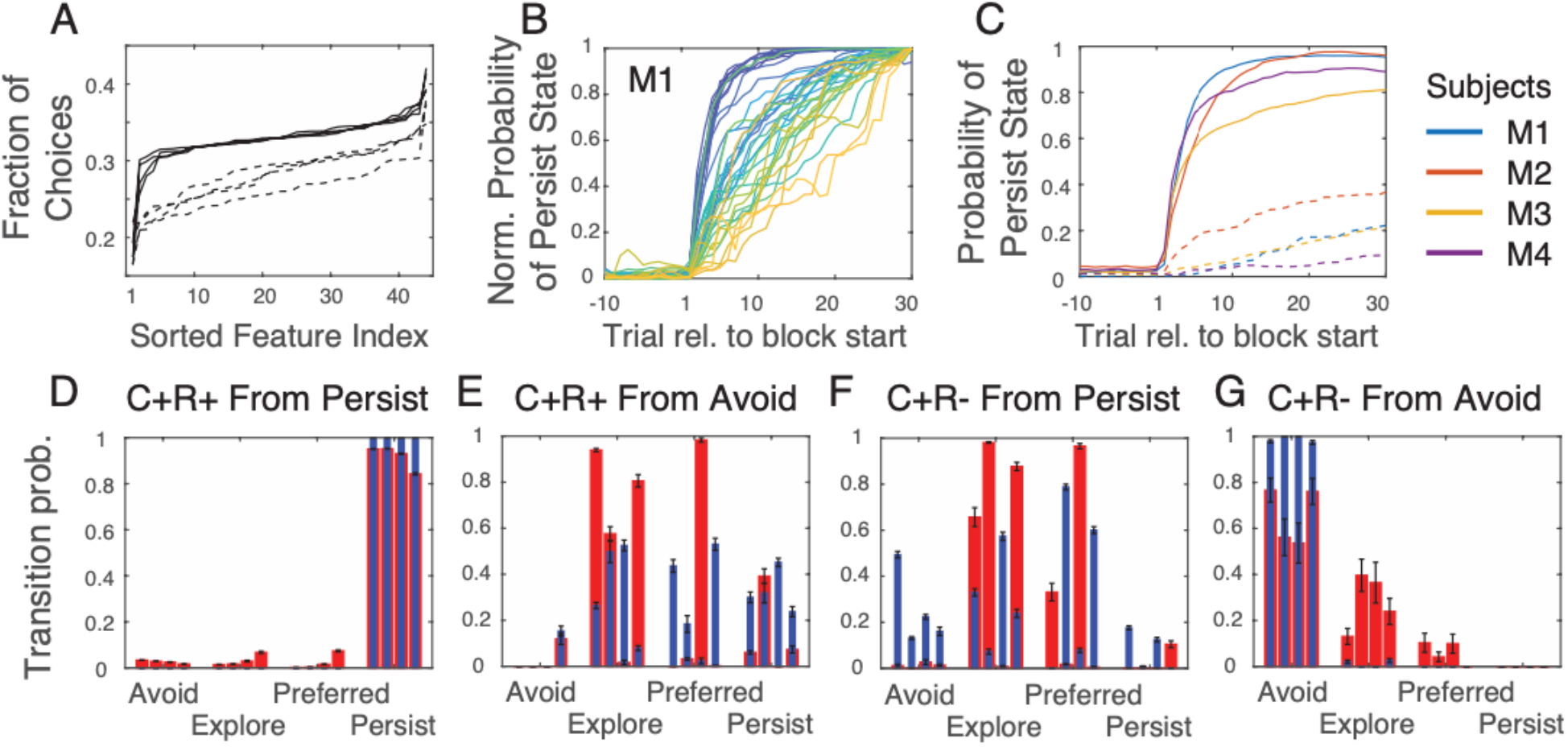
State transitions depend on long-term feature preference. (A) Fraction of trials on which a particular feature is present in the chosen object (solid lines) across all trials and (dashed line) across trials in which it was not the rewarded feature, ordered from low (less preferred) to high (more preferred). Each curve represents a separate subject. (B) Norm. probability of the target feature-channel being in the Persist state in trials relative to block start, with a curve for each feature, color coded from blue to yellow for most preferred to least preferred feature (ordered based on fraction chosen when not the target feature). (C) Mean across (solid) most preferred ten and (dashed) least preferred seven for each subject, color coded as indicated in the legend. (D-G) Transition matrices obtained for ten most preferred (red, wide bars) versus seven least preferred (blue thin bars) features for each of the 4 subjects, shown as bars stacked next to each other. Transitions are shown when the feature was rewarded and chosen in the previous trial (C+R+) and the feature-channel was in the Persist state (D) or in the Avoid state (E) and for trials after a non-rewarded choice of an object with that feature (C+R-) when the feature-channel was in the Persist state (F) or in the Avoid state (G). When a choice led to a reward, the state mostly remained Persist, whereas the state transitions away from Avoid. When a feature choice did not lead to a reward, states transition away from Persist, with non-preferred features moved more to the Avoid state than the preferred feature. Non-preferred features also mostly stayed in Avoid, whereas a sizeable fraction of states of preferred features moved towards the Explore state.

## References

Allen, R.J. & Ueno, T. (2018) Multiple high-reward items can be prioritized in working memory but with greater vulnerability to interference. Atten Percept Psychophys, 80, 1731–1743.

Anderson, B.A., Kim, H., Kim, A.J., Liao, M.R., Mrkonja, L., Clement, A. & Gregoire, L. (2021) The past, present, and future of selection history. Neurosci Biobehav R, 130, 326–350.

Ashwood, Z.C., Roy, N.A., Stone, I.R., Urai, A.E., Churchland, A.K., Pouget, A., Pillow, J.W. & Lab, I.B. (2022) Mice alternate between discrete strategies during perceptual decision-making. Nat Neurosci, 25, 201-+.

Awh, E., Belopolsky, A.V. & Theeuwes, J. (2012) Top-down versus bottom-up attentional control: a failed theoretical dichotomy. Trends Cogn Sci, 16, 437–443.

Baddeley, A. (2012) Working Memory: Theories, Models, and Controversies. Annu Rev Psychol, 63, 1–29.

Bengio, Y. & Frasconi, P. (1994) An Input Output HMM Architecture. In Tesauro, G., Touretzky, D., Leen, T. (eds).

Boroujeni, K.B., Sigona, M.K., Treuting, R.L., Manuel, T.J., Caskey, C.F. & Womelsdorf, T. (2022a) Anterior cingulate cortex causally supports flexible learning under motivationally challenging and cognitively demanding conditions. Plos Biol, 20.

Boroujeni, K.B., Watson, M. & Womelsdorf, T. (2022b) Gains and Losses Affect Learning Differentially at Low and High Attentional Load. J Cognitive Neurosci, 34, 1952–1971.

Collins, A.G.E. & Frank, M.J. (2012) How much of reinforcement learning is working memory, not reinforcement learning? A behavioral, computational, and neurogenetic analysis. Eur J Neurosci, 35, 1024–1035.

Cowan, N., Bao, C., Bishop-Chrzanowski, B.M., Costa, A.N., Greene, N.R., Guitard, D., Li, C., Musich, M.L. & Unal, Z.E. (2024) The Relation Between Attention and Memory. Annu Rev Psychol, 75, 183–214.

Dempster, A.P., Laird, N.M. & Rubin, D.B. (1977) Maximum Likelihood from Incomplete Data Via Em Algorithm. J Roy Stat Soc B Met, 39, 1–38.

Farashahi, S., Rowe, K., Aslami, Z., Lee, D. & Soltani, A. (2017) Feature-based learning improves adaptability without compromising precision. Nat Commun, 8.

Frémaux, N. & Gerstner, W. (2016) Neuromodulated Spike-Timing-Dependent Plasticity, and Theory of Three-Factor Learning Rules. Front Neural Circuit, 9.

Ghazizadeh, A., Griggs, W. & Hikosaka, O. (2016) Ecological Origins of Object Salience: Reward, Uncertainty, Aversiveness, and Novelty. Front Neurosci-Switz, 10.

Goudar, V., Kim, J.W., Liu, Y., Dede, A.J.O., Jutras, M.J., Skelin, I., Ruvalcaba, M., Chang, W.L., Ram, B., Fairhall, A.L., Lin, J.J., Knight, R.T., Buffalo, E.A. & Wang, X.J. (2024) A Comparison of Rapid Rule-Learning Strategies in Humans and Monkeys. J Neurosci, 44.

Goudar, V., Peysakhovich, B., Freedman, D.J., Buffalo, E.A. & Wang, X.J. (2023) Schema formation in a neural population subspace underlies learning-to-learn in flexible sensorimotor problem-solving. Nat Neurosci, 26, 879-+.

Hanks, T.D. & Summerfield, C. (2017) Perceptual Decision Making in Rodents, Monkeys, and Humans. Neuron, 93, 15–31.

Hitch, G.J., Hu, Y., Allen, R.J. & Baddeley, A.D. (2018) Competition for the focus of attention in visual working memory: perceptual recency versus executive control. Ann N Y Acad Sci, 1424, 64–75.

Jahn, C.I., Markov, N.T., Morea, B., Daw, N.D., Ebitz, R.B. & Buschman, T.J. (2024) Learning attentional templates for value-based decision-making. Cell, 187.

Johnston, W.J. & Fusi, S. (2023) Abstract representations emerge naturally in neural networks trained to perform multiple tasks. Nat Commun, 14.

Kemp, C. & Tenenbaum, J.B. (2008) The discovery of structural form. P Natl Acad Sci USA, 105, 10687–10692.

Khamassi, M., Wilson, C.R.E., Rothé, M., Quilodran, R., Dominey, P.F. & Procyk, E. (2011) Meta-Learning, Cognitive Control, and Physiological Interactions between Medial and Lateral Prefrontal Cortex. Neural Basis of Motivational and Cognitive Control, 351–369.

Kobak, D., Brendel, W., Constantinidis, C., Feierstein, C.E., Kepecs, A., Mainen, Z.F., Qi, X.L., Romo, R., Uchida, N. & Machens, C.K. (2016) Demixed principal component analysis of neural population data. Elife, 5.

Kriegeskorte, N. & Kievit, R.A. (2013) Representational geometry: integrating cognition, computation, and the brain. Trends Cogn Sci, 17, 401–412.

Leong, Y.C., Radulescu, A., Daniel, R., DeWoskin, V. & Niv, Y. (2017) Dynamic Interaction between Reinforcement Learning and Attention in Multidimensional Environments. Neuron, 93, 451–463.

Mansouri, F.A., Buckley, M.J., Fehring, D.J. & Tanaka, K. (2020) The Role of Primate Prefrontal Cortex in Bias and Shift Between Visual Dimensions. Cereb Cortex, 30, 85–99.

Miller, E.K. & Buschman, T.J. (2013) Cortical circuits for the control of attention. Curr Opin Neurobiol, 23, 216–222.

Oemisch, M., Westendorff, S., Azimi, M., Hassani, S.A., Ardid, S., Tiesinga, P. & Womelsdorf, T. (2019) Feature-specific prediction errors and surprise across macaque fronto-striatal circuits. Nat Commun, 10.

Ostojic, S. & Fusi, S. (2024) Computational role of structure in neural activity and connectivity. Trends Cogn Sci, 28.

Pearson, D., Chong, A., Chow, J.Y.L., Garner, K.G., Theeuwes, J. & Le Pelley, M.E. (2024) Uncertainty-Modulated Attentional Capture: Outcome Variance Increases Attentional Priority. J Exp Psychol Gen, 153, 1628–1643.

Pearson, D. & Le Pelley, M.E. (2020) Learning to avoid looking: Competing influences of reward on overt attentional selection. Psychon B Rev, 27, 998–1005.

Rusz, D., Le Pelley, M.E., Kompier, M.A.J., Mait, L. & Bijleveld, E. (2020) Reward-Driven Distraction: A Meta-Analysis. Psychol Bull, 146, 872–899.

Rutishauser, U., Slotine, J.J. & Douglas, R.J. (2012) Competition Through Selective Inhibitory Synchrony. Neural Comput, 24, 2033–2052.

Shadlen, M.N. & Kiani, R. (2013) Decision Making as a Window on Cognition. Neuron, 80, 791–806.

Smith, A.C., Frank, L.M., Wirth, S., Yanike, M., Hu, D., Kubota, Y., Graybiel, A.M., Suzuki, W.A. & Brown, E.N. (2004) Dynamic analysis of learning in behavioral experiments. J Neurosci, 24, 447–461.

Soto, F.A., Vogel, E.H., Uribe-Bahamonde, Y.E. & Perez, O.D. (2023) Why is the Rescorla-Wagner model so influential? Neurobiol Learn Mem, 204.

Souza, A.S. & Oberauer, K. (2016) In search of the focus of attention in working memory: 13 years of the retro-cue effect. Atten Percept Psychophys, 78, 1839–1860.

Sutton, R.S. & Barto, A.G. (2018) Reinforcement learning : an introduction. The MIT Press, Cambridge, Massachusetts.

Tian, Z.H., Chen, J.W., Zhang, C., Min, B., Xu, B. & Wang, L.P. (2024) Mental programming of spatial sequences in working memory in the macaque frontal cortex. Science, 385.

Treuting, R.L., Banaie Boroujeni, K., Gerrity, C.G., Neumann, A., Tiesinga, P. & Womelsdorf, T. (2025) Adaptive reinforcement learning is causally supported by anterior cingulate cortex and striatum. Neuron.

Usher, M. & Niebur, E. (1996) Modeling the temporal dynamics of IT neurons in visual search: A mechanism for top-down selective attention. J Cognitive Neurosci, 8, 311–327.

Wang, J.X., Kurth-Nelson, Z., Kumaran, D., Tirumala, D., Soyer, H., Leibo, J.Z., Hassabis, D. & Botvinick, M. (2018) Prefrontal cortex as a meta-reinforcement learning system. Nat Neurosci, 21, 860-+.

Watson, M.R., Traczewski, N., Dunghana, S., Boroujeni, K.B., Neumann, A., Wen, X. & Womelsdorf, T. (2023) A Multi-task Platform for Profiling Cognitive and Motivational Constructs in Humans and Nonhuman Primates. bioRxiv, 2023.2011.2009.566422.

Watson, M.R., Voloh, B., Naghizadeh, M. & Womelsdorf, T. (2019) Quaddles: A multidimensional 3-D object set with parametrically controlled and customizable features. Behav Res Methods, 51, 2522–2532.

Whittington, J.C.R., McCaffary, D., Bakermans, J.J.W. & Behrens, T.E.J. (2022) How to build a cognitive map. Nat Neurosci, 25, 1257–1272.

Williams, A.H., Kim, T.H., Wang, F., Vyas, S., Ryu, S.I., Shenoy, K.V., Schnitzer, M., Kolda, T.G. & Ganguli, S. (2018) Unsupervised Discovery of Demixed, Low-Dimensional Neural Dynamics across Multiple Timescales through Tensor Component Analysis. Neuron, 98, 1099-+.

Williamson, R.C., Doiron, B., Smith, M.A. & Yu, B.M. (2019) Bridging large-scale neuronal recordings and large-scale network models using dimensionality reduction. Curr Opin Neurobiol, 55, 40–47.

Womelsdorf, T., Thomas, C., Neumann, A., Watson, M.R., Boroujeni, K.B., Hassani, S.A., Parker, J. & Hoffman, K.L. (2021) A Kiosk Station for the Assessment of Multiple Cognitive Domains and Cognitive Enrichment of Monkeys. Front Behav Neurosci, 15.

Womelsdorf, T., Watson, M.R. & Tiesinga, P. (2022) Learning at Variable Attentional Load Requires Cooperation of Working Memory, Meta-learning, and Attention-augmented Reinforcement Learning. J Cognitive Neurosci, 34, 79–107.

